# In utero Bisphenol A exposure developmentally reprograms mammary gland fibroblasts

**DOI:** 10.64898/2026.07.08.737306

**Authors:** Jillian M. Poska, Bader Albalawi, Madeline R. Price, Craig J. Burd

## Abstract

In utero exposure to estrogens is known to increase breast cancer risk in adulthood. The plasticizer bisphenol A (BPA) increases tumor susceptibility in rodent models of in utero exposure, but the mechanism of this increase is unclear. Our lab has previously shown that in utero BPA exposure causes alterations to the tissue microenvironment that are conducive to tumor initiation. We thus sought to understand when these alterations arise and how they are regulated using bulk and single-nuclei epigenomics in adult mammary glands following in utero BPA exposure. Our results indicate that in utero BPA exposure causes developmental reprogramming of mammary fibroblasts that is initiated during puberty and persists into adulthood. Fibroblast-specific changes in chromatin accessibility were identified at enhancers and annotated to genes involved in fibroblast differentiation, suggesting that BPA may exert its effects by reprogramming fibroblast heterogeneity. Single-nuclei multiomics confirmed changes in fibroblast heterogeneity and uncovered estrogen signaling as a putative regulator of the fibroblast cell state that is altered by BPA exposure. Together, these studies support a model whereby BPA acts on mesenchymal ERα to reprogram the chromatin landscape of mammary fibroblasts, leading to their altered differentiation.

## Introduction

Though estrogen signaling is critical for mammary gland development, inappropriate exposure to estrogens can alter developmental processes and increase breast cancer risk. This was first evidenced by diethylstilbestrol (DES), which increased breast cancer risk twofold after age 40 in women who were exposed in utero^1^. Though DES is no longer prescribed, humans are frequently exposed to other chemicals that affect hormone signaling, termed endocrine disrupting compounds (EDCs). One common EDC is bisphenol A (BPA), a compound that is found in plastics, can linings, dental sealants, and thermal receipt paper and is known to engage nuclear hormone receptors, including estrogen receptor alpha (ERα)^2^. BPA can be detected in human amniotic fluid, raising the concern that in utero exposure can increase breast cancer risk similarly to DES^3,4^. Rodents exposed in utero to BPA undergo altered mammary epithelial development and have increased mammary tumor susceptibility following carcinogenic challenge^5–12^. However, it is still not fully understood how BPA exerts this effect, making it difficult to assess the risk of exposure and design alternatives that avoid pathways of concern.

The developmental origins of health and disease (DOHaD) theory has been hypothesized to explain how exposure to BPA in utero can increase breast cancer risk in adulthood. This theory posits that the fetal environment is critical for instructing tissue development and thus disruptions to this environment have the potential to increase disease risk in adulthood^13^. In the mouse, fetal mammary gland development begins with the formation of epithelial buds from the overlying ectoderm, which are surrounded by a dense mesenchymal layer^14^. The buds sprout and invade the fat pad precursor in the later stages of embryonic development, then grow allometrically until puberty, when the increase in circulating hormones causes rapid ductal expansion and proliferation^15^. The ductal structure is comprised of both luminal and basal epithelial cells. The epithelium is embedded in an adipose-rich stroma, comprised of adipocytes, immune cells, and mesenchyme-derived fibroblasts, which provide cues for growth, differentiation, and function of the mammary gland^16^. This cellular complexity makes it challenging to study developmental reprogramming as it is difficult to assess the contribution of each cell type to the exposure phenotype.

To assess the effect of in utero BPA exposure in individual cell types of the mammary gland, our lab previously performed RNA-seq on isolated luminal, basal, and fibroblast cells from adult mice exposed in utero to BPA. Differentially expressed genes in fibroblasts were associated with breast cancer pathways; specifically, BPA-exposed fibroblasts upregulated multiple collagens^17^. Functionally, BPA exposure increased collagen deposition and mammary tissue stiffness^17,18^, which are known risk factors for breast cancer^19,20^. This data suggested that BPA could increase breast cancer risk by promoting a mammary microenvironment that promotes tumorigenesis. Supporting this hypothesis, mesenchymal knockout of ERα protects mice from tumor susceptibility driven by in utero BPA exposure^21^. RNA-seq of tissue compartments in exposed embryos additionally shows that the mesenchyme is transcriptionally altered by BPA at the time of exposure^22^. However, pre-pubertal mice exposed in utero to BPA do not show changes in collagen deposition or tissue stiffness, suggesting that the changes initiated in utero by BPA are remembered later in life. The mechanism by which memory of BPA exposure is propagated to adulthood is presently unclear.

Memory is often associated with epigenetic changes that prime genes to change their response to stimulus^23^. Epigenetic mechanisms such as changes in DNA methylation and histone modification have been linked to adverse effects of developmental exposure to pollutants, including cigarette smoke and heavy metals^24,25^. In utero BPA exposure has previously been shown to alter DNA methylation across the genome and increase H3K4me3 at the α-lactalbumin promoter^26^. Additionally, mice exposed in utero to BPA upregulated the enzyme EZH2 in the mammary gland and showed a corresponding increase in H3K27me3, though the targets of this modification are not known^27^. However, these studies were performed in whole mammary tissue, obscuring the cellular targets of epigenetic reprogramming.

We thus aimed to elucidate the mechanism of developmental reprogramming by BPA in the mammary gland, specifically focusing on the mammary stroma. We analyzed primary mammary gland cell types from mice exposed in utero to BPA to identify transcriptional and epigenetic changes that occur following BPA exposure. The results of these experiments add further understanding to the mechanism of BPA exposure in increasing cancer risk and uncover fibroblast differentiation as a novel endpoint affected by estrogen exposure.

## Methods

### Animals

Animal experiments were performed in compliance with protocols approved by The Ohio State University Institutional Animal Care and Use Committee (IACUC, Protocol #2013A00000030) and in accordance with the accepted standard of humane animal care. CD1 mice (Charles River, Wilmington, MA, USA) were maintained in polysulfone cages and fed a diet containing minimal levels of phytoestrogen (Harlan Teklad 2919). Sexually mature female CD1 mice (aged ≥ 8 weeks) were mated, and appearance of a vaginal plug was taken to be embryonic day (E) 0.5, at which point they were randomly assigned to a treatment group. BPA (Sigma 235698, ≥99%) dissolved in sesame oil was administered to pregnant mice via oral gavage. Treatment of pregnant dams with either 25 µg/kg body weight (bw)/day of BPA or an equivalent dose of sesame oil occurred daily from E9.5 through E18.5. The 25 µg/kg bw dose was previously shown to lead to amniotic fluid concentrations of BPA similar to those reported in humans^12^. For all tissue collection, mice were euthanized by CO2 inhalation and euthanasia was confirmed by cervical dislocation.

### Isolation of primary cells

Fourth and fifth inguinal mammary glands were removed from adult female mice at the timepoints indicated for each experiment. Primary cells from the mammary gland were isolated and sorted as previously described^28^. Briefly, glands with the lymph nodes removed were mechanically minced and digested in a mixture of collagenase A and trypsin. Cells underwent red blood cell lysis and washing, after which fibroblasts were separated by adhesion to a cell culture dish. The non-adherent cells were further processed to a single-cell suspension by trypsinization and DNAse I treatment and counted. Processed cells were stained for cell sorting using antibodies against CD24 (BD 553261, 1:80), CD49f (BD 551129 1:8), and CD45 (BD 552848, 1:50). CD45+ cells were removed and epithelial cell populations were separated by their expression of CD24 and CD49f.

### qRT-PCR

DNA and RNA were isolated from fibroblasts using the Quick DNA/RNA mini prep kit (Zymo Research) the morning after fibroblasts were isolated. 1 µg of RNA was converted to cDNA using the iScript Select cDNA Synthesis Kit (BioRad, 170-8897). Relative gene expression was measured by qPCR with Forget-Me-Not Evagreen SYBR (VWR, 76204) using previously reported collagen primers, with *Rpl13a* as a control^17^.

### Picrosirius red staining

Fourth inguinal mammary glands were fixed for 72 hours in 10% NBF, after which they were embedded in paraffin wax and sectioned. Picrosirius red staining was performed as previously described (22,23). Briefly, sections were rehydrated through a series of alcohols and incubated for 1 hour in picrosirius red stain (Abcam 68742), after which they were rinsed in two changes of 0.5% acetic acid and mounted. To quantify staining, two images were taken per section adjacent to the lymph node. Images were adjusted to remove any portions of the lymph node, tissue tears, and blank areas of the slide, then scored in ImageJ by measuring the area with signal above a standardized threshold on the red channel. Data is displayed as a percentage of area meeting the threshold over total area in the image.

### ATAC-seq

After cell sorting, 50,000 primary cells of each cell type were aliquoted for assay for transposase accessible chromatin and sequencing (ATAC-seq). Nuclei were isolated, DNA was tagmented, and libraries were prepared using the Diagenode ATAC-seq kit (Cat. C01080002). Size distribution was assessed by Tapestation HS1000 before sequencing.

Reads were deduplicated using fastuniq and adapter sequences were removed using cutadapt^29,30^. Paired-end reads were aligned to the mm10 genome using Bowtie2 and samtools was used to filter unmapped reads, reads failing platform QC, and improperly aligned reads^31,32^.

Peaks were called using HMMRATAC on MACS3 and regions mapping to the ENCODE blacklist were removed^33–35^. Peak calls for each sample within a given cell type were merged using HOMER to create a union peak list of accessible chromatin in each cell type^36^. For each cell type, reads were counted within peaks using featureCounts^37^. Differential accessibility was assessed from these counts using the EdgeR^38^ framework, with │log2FC│ > 1.5 and FDR < 0.05.

Positioning of differentially accessible regions (DARs) from transcription start sites was assessed using the Genomic Regions Enrichment of Annotations Tool^39^. DARs were annotated and assessed for the enrichment of transcription factor motifs using HOMER AnnotatePeaks and FindMotifsGenome, respectively. ToppFun was used to identify pathways enriched for both nearest genes and enriched transcription factors^40^. Nearest genes to DARs were additionally overlapped with fibroblast subtype markers from a published single cell dataset of mammary fibroblasts^41^. As a control comparison, cell type specific genes were overlapped with 3 independent random gene sets generated using the random gene set generator from MolBioTools.com.

### CUT&Tag

After cell isolation, 100,000 primary fibroblasts were aliquoted for cleavage under targets and tagmentation (CUT&Tag). CUT&Tag was performed as previously described with antibodies targeting H3K4me1 (Cell Signaling 5326, 1:100) or IgG (Cell Signaling 2729, 1:100)^42^. Size distribution was assessed by Tapestation HS1000 before sequencing.

Reads were processed and aligned to the mm10 genome as described for ATAC-seq. Peaks were called against the IgG control for each treatment group separately using MACS3 callpeak with the broad setting. Peak calls in each treatment group were merged using HOMER mergePeaks and reads were counted in the union peak list for each sample using featureCounts. Differential methylation was assessed from these counts using EdgeR with │log2FC│ > 1.5 and FDR < 0.05. Peaks were overlapped with DARs using HOMER mergePeaks.

### Sample preparation for single nuclei multiome sequencing

Mammary glands were processed as described above excluding the fibroblast adhesion step. After red blood cell lysis, cell pellets were dissociated to single cell suspensions and cryopreserved until nuclei isolation.

Nuclei were isolated following 10X Genomics protocol CG000365. Lysis buffer was used at a 0.1X dilution as recommended for primary cells and lysis was carried out for 3 minutes. Nuclei were counted using DAPI and diluted to the appropriate concentration before sending to the Ohio State University Genomics Shared Resource for barcoding and library preparation.

### Multiome sequencing alignment and quality control

Code used to analyze the single cell data can be found at (https://github.com/poskajm/Developmental-Reprogramming-Paper-Code). Sequencing libraries were aligned to the mm10 genome, and a feature barcode matrix was made for each sample using CellRanger ARC v2.0.2. Seurat v5.2.0 and Signac v1.16.0 were used to analyze gene expression (GEX) and chromatin accessibility (ATAC) modalities^43,44^. Seurat objects for each treatment were merged to generate a single object for each timepoint.

Quality control was performed on each sample individually before merging. UMI counts were adjusted for ambient RNA contamination using SoupX^45^. The top ambient genes as determined by SoupX were additionally used as input in the PercentageFeatureSet() function to filter cells with high ( > 5%) contributions from the ambient RNA. Cells with outlier read counts, as determined for each sample by density scatter plots, were filtered out. Additionally, cells with high nucleosomal signal ( > 2), low transcription start site (TSS) enrichment ( < 3) or high percentage of mitochondrial genes (> 20) were removed.

### Multiome normalization and clustering

Downstream analysis was performed on the merged dataset including both BPA and vehicle treated samples. GEX data was normalized and scaled using variance stabilizing transformation (VST) and integrated using Harmony. ATAC data was normalized using latent semantic indexing (LSI) and integrated with the mutual nearest neighbor approach using the FindIntegrationAnchors() function of Seurat. Dimensional reduction was performed using a weighted nearest neighbor (WNN) approach considering both GEX and ATAC modalities followed by UMAP visualization. Clusters were identified from the WNN UMAP using the SLM algorithm with resolution = 0.2.

The FindAllMarkers() function in Seurat was used to identify both genes and accessible regions enriched in each cluster. RNA markers were considered to define a cluster if they were expressed in at least 30% of cells in that cluster and had a logFC ≥ 0.5 and padj ≤ 0.05. ATAC regions were determined by expression in at least 10% of that cluster with an adjusted p value of ≤ 0.05. RNA markers were manually compared to published single cell gene expression data sets from adult mammary and inguinal adipose tissue in order to determine cluster identity^46,47^. Module scores were calculated for the Pi16 signature based on a pan-tissue fibroblast data set^48^.

### Downstream analysis of multiome data

Differential gene expression was performed on each cluster using the MAST algorithm, which has previously been suggested to the best performing differential expression test for single samples^49^. Differential expression was defined with a logFC threshold of 0.5 and an adjusted p value of < 0.05. Additionally differentially expressed genes (DEGs) were filtered to those expressed in at least 20% of one sample to avoid biasing lowly expressed genes. Lists of DEGs were filtered against lists of genes highly expressed in the ambient RNA to avoid detecting differences due to sample quality. Differential gene expression analysis was repeated with the same parameters on groups of clusters from the same cell type (all fibroblasts, all luminal cells, and all basal cells) to compare with previously published bulk RNA-seq data^17^. Differential accessibility of ATAC peaks was assessed with a log-rank test using the same logFC and adjusted p value thresholds, but with a minimum percent expression of 10%.

Fibroblasts were subset and reclustered using the WNN algorithm. Markers were found for each subcluster as described above. A module score was added for Pi16 fibroblasts using a previously published gene signature of this cell type^48^. Fibroblast trajectory was predicted using Slingshot using Pi16+ fibroblasts as a starting cluster^50^. ATAC markers for fibroblast subtypes were overlapped with DARs from bulk ATAC data using HOMER mergePeaks.

Motif accessibility was assessed with chromVAR using the mouse_pwm_v2 motif set from the chromVARmotifs package^51^. Differences in motif accessibility were assessed with a Wilcoxon rank-sum test, with differentially enriched motifs having a logFC threshold of 0.5 and p_val_adj < 0.05. Gene regulatory networks (GRNs) were reconstructed for fibroblasts using Pando, which leverages both the ATAC and RNA modalities to assess gene expression and coordinating motif accessibility in the same cell^52^. The identified GRNs were added as module scores to the Seurat object and GRN enrichment was assessed with the same parameters as described for chromVAR.

### Flow cytometry

Adult mammary glands were processed to single cell suspensions as described for single nuclei multiome sequencing. Cells were counted and resuspended in FACS buffer at a concentration of 1×10^6^ cells/100 µL. Adult cells were stained with fluorescently conjugated antibodies against CD45 (Tonbo 50-0451-U100, 1:300), CD31 (Biolegend 102526, 1:40), CD24 (Biolegend 101827, 1:100), CD49f (BD 551129, 1:8), DPP4 (Biolegend 137810, 1:100), CD55 (Biolegend 131811, 1:200), and CD9 (Biolegend 124808, 1:100). Percentage of DPP4 positivity within stromal fibroblasts (CD45^-^CD31^-^CD49f^-^CD24^-^) was compared between BPA and vehicle treated mammary glands with an unpaired t-test.

## Results

### Collagen expression is developmentally controlled

We previously showed that the collagen deposition and mammary gland stiffness phenotypes of in utero BPA exposure are not evident in exposed mice at 4 weeks of age^17^. In order to assess if this observation is controlled at the transcriptional level, we performed qRT-PCR for collagen genes in mammary fibroblasts from 4 week old mice exposed in utero to BPA or a vehicle control. There were no significant differences in expression of any of the tested collagens between BPA and vehicle treated mice (Figure 1A), confirming our prior tissue-level data. To narrow down the time between puberty and adulthood when collagen is upregulated in BPA-exposed mice, we additionally performed qRT-PCR at 7 and 8 weeks of age (Figure 1B,C). There were no significant differences seen at 7 weeks, but there were several transcripts that begin to demonstrate a trend towards increased mRNA expression in BPA-exposed mice compared to vehicle controls (Col1a1, Col1a2, Col3a1, Col4a1, Col6a3, Col6a2) (Figure 1B). At 8 weeks, BPA significantly increased the expression of Col1a2 and Col4a1 (Figure 1C). Though not all statistically significant, most of the tested collagens also showed higher expression in BPA-exposed mice at this timepoint.

**Figure 1.**
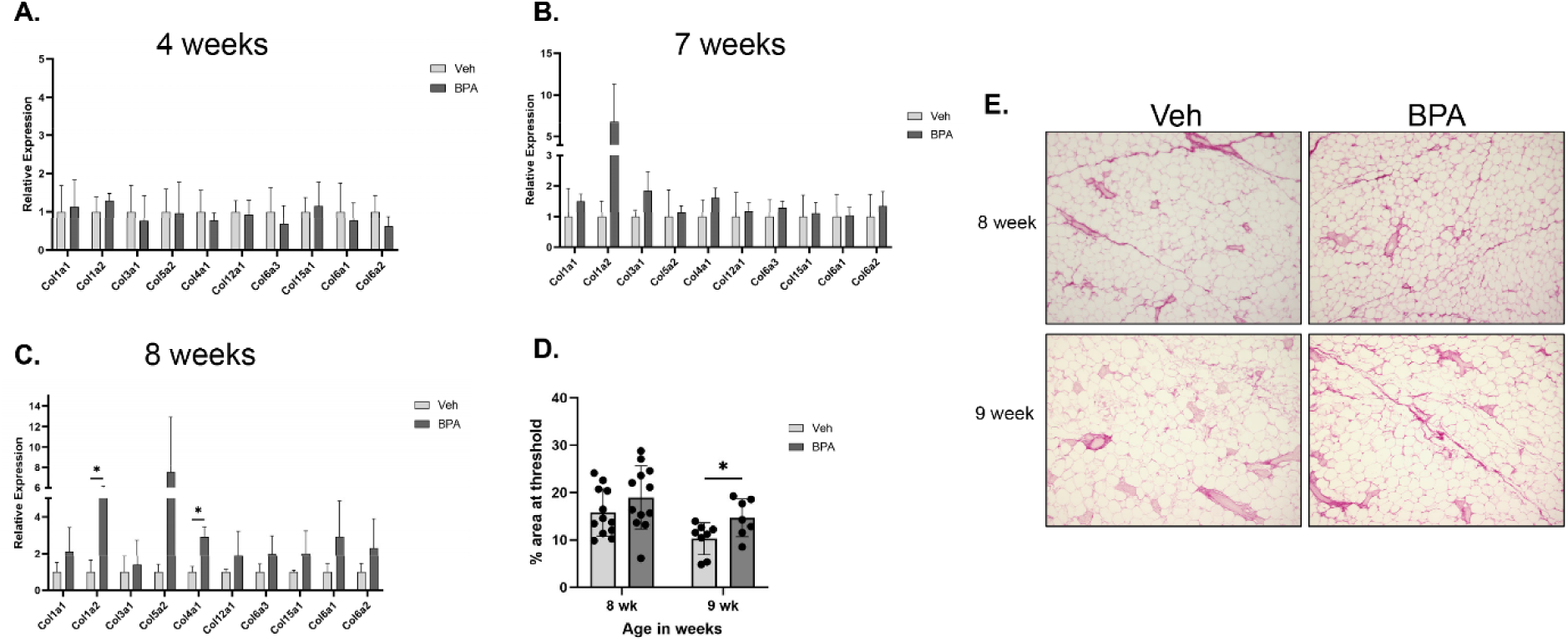
In utero BPA results in increased collagen expression during mammary gland maturation. RNA was isolated from fibroblasts at the indicated timepoints and used for qRT-PCR. Ct values were normalized to Rpl13a to determine expression at 4 weeks (A), 7 weeks (B), or 8 weeks (C) of age after in utero BPA exposure. Plots indicate the relative expression as normalized to vehicle treated samples (n = 3). Collagen expression was validated by staining sectioned tissue with picrosirius red and measuring the percentage of pixels above a red threshold (D). Each point represents the average of two images taken at 4x magnification for one mouse. Representative images of picrosirius red staining are shown (E).

In order to determine if the transcriptional increases in collagen were reflected in increased collagen deposition at 8 weeks, mammary tissue sections were stained with picrosirius red, which marks fibrillar collagen. At 8 weeks of age, BPA exposed mice showed a trend of increased collagen deposition, but this change was not statistically significant. When we extended this analysis to 9 weeks, we were able to detect a statistically significant difference between BPA and vehicle treated mice (Figure 1D), which can be visualized in representative images of mice at each timepoint (Figure 1E). Interestingly, both groups showed lower collagen deposition at 9 weeks compared to 8 weeks, but BPA-treated mice had a smaller magnitude of decrease. Altogether, these data suggest that BPA exposure causes dysregulation of normal mammary stromal development that manifests during mammary gland maturation.

### BPA exposure alters chromatin accessibility at fibroblast enhancers

The transcriptional changes in collagen that occur much later than the initial exposure suggest a mechanism of transcriptional memory regulating the exposure phenotype. In order to determine the mechanism and cell-type specificity of epigenetic regulation following in utero BPA exposure, we performed assay for transposase accessible chromatin sequencing (ATAC-seq) on sorted luminal epithelial cells, basal epithelial cells, and fibroblasts at 12 weeks of age, a timepoint that previously showed transcriptional dysregulation and changes in the tissue microenvironment. Differentially accessible regions (DARs) were determined using EdgeR with the criteria of │FC│ ≥ 1.5 and FDR ≤ 0.05, which revealed very few differences in chromatin accessibility in either epithelial population. In contrast, fibroblasts showed 384 regions with enriched accessibility in vehicle and 183 regions with enriched accessibility in BPA treatment (Figure 2A).

**Figure 2.**
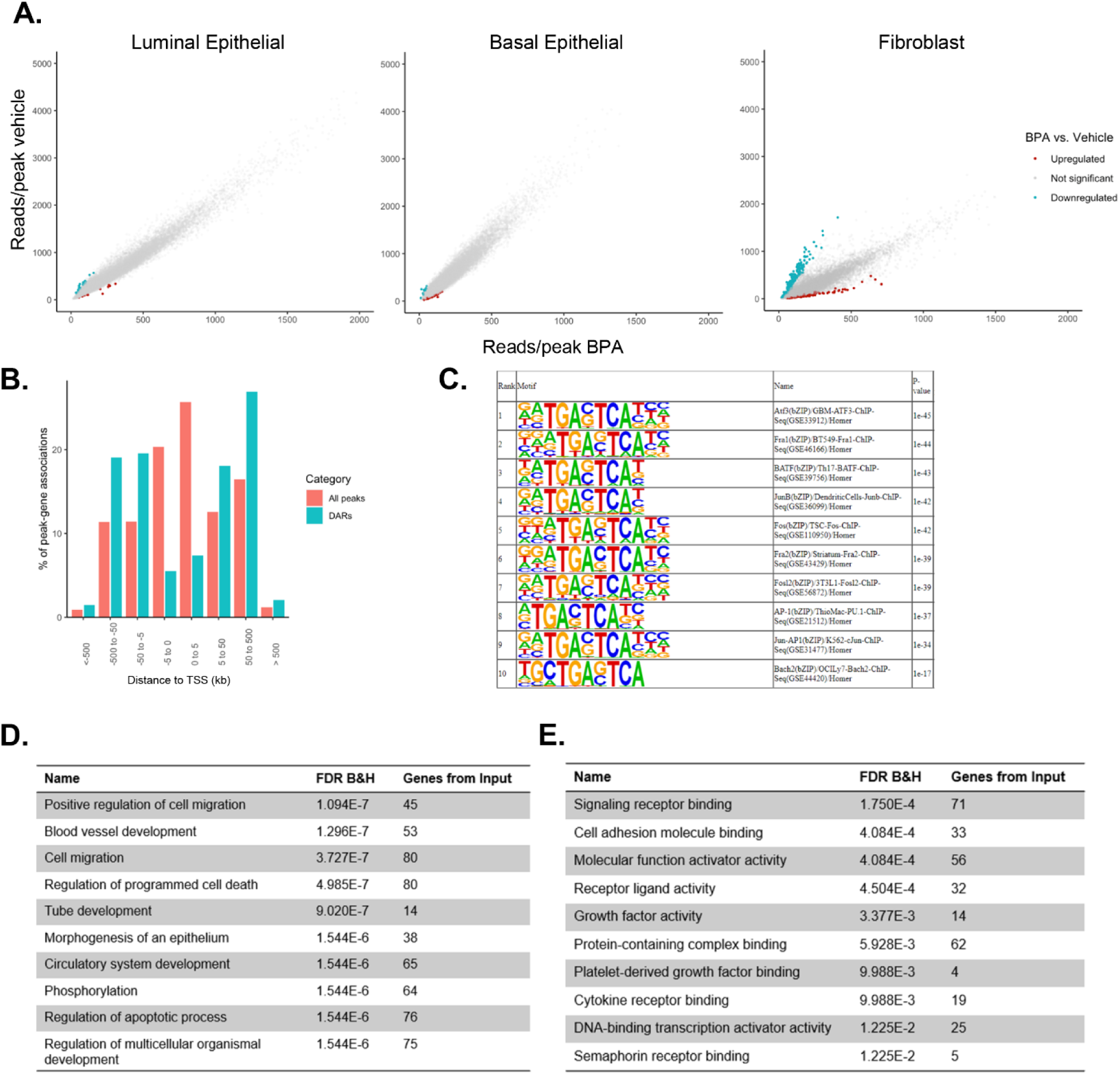
The fibroblast chromatin landscape is reprogrammed by in utero BPA exposure. Cell populations isolated by FACS were subjected to ATAC-seq library preparation using the Diagenode kit. Sequencing reads were aligned to the mm10 genome and peaks were called using HMMRATAC. Normalized reads per peak were averaged within treatment groups (n=3) and plotted as counts in vehicle treated mice vs. BPA treated mice. Differentially accessible region for BPA (red) or vehicle (blue) treatment were defined as peaks where │log2FC│ ≥ 1.5 and FDR≤ 0.05 **(A)**. Genomic Regions Enrichment of Annotations (GREAT) was used to annotate peaks in fibroblasts. Percent of peak-gene associations is plotted against distance from that gene’s TSS for all peaks (pink) and DARs (blue) **(B)**. HOMER was used to determine transcription factor motifs enriched in differentially accessible regions. The top 10 motifs in fibroblasts are shown **(C)**. The nearest genes to each DAR were used as the input for gene ontology analysis using the ToppGene suite. The top biological process **(D)** and molecular function **(E)** terms are shown.

Accessible peaks in fibroblasts were enriched within 5kb of transcription start sites (TSSs), demonstrating that the ATAC-seq protocol correctly profiled regions that are naturally more accessible (Figure 2B, pink bars). In contrast, DARs in fibroblasts were enriched in regions further than 5 kb from the nearest transcription start site, suggesting that BPA exposure primarily affects distal regulatory elements (Figure 2B, blue bars). Fibroblast DARs showed enrichment of transcription factor motifs from the AP-1 family, which are known to tether ERα and have been associated with enhancers that change between normal mammary fibroblasts and cancer associated fibroblasts (Figure 2C)^53,54^. DARs were further annotated using HOMER and the closest genes were analyzed for enriched pathways using the ToppGene suite^40^. The top biological processes enriched in DARs included terms related to migration and development (Figure 2D) and the top molecular function terms included signaling receptor binding, cell adhesion molecule binding, and growth factor activity (Figure 2E). Importantly, we did not identify any significantly enriched terms relating to extracellular matrix production or organization, indicating that BPA may not be acting directly on collagen genes. Altogether, these data suggest that BPA exposure reprograms distal regulatory elements in fibroblasts to affect cell motility, morphogenesis, and communication.

Because DARs in fibroblasts were found to be enriched in regions distal to TSSs, we performed CUT&Tag for H3K4me1 to examine whether BPA exposure affected enhancers in the fibroblast genome. 314 out of 556 DARs overlapped with H3K4me1 peaks, suggesting that BPA exposure affects chromatin accessibility primarily at enhancers (Figure 3A). However, analysis of H3K4me1 counts within DARs revealed very few significant changes in H3K4me1 deposition with BPA exposure (Figure 3B). Additionally, minimal changes were identified in H3K4me1 counts within called peaks (Figure 3C). Further validating this finding, very few DARs overlapped with H3K4me1 peaks that were called uniquely in vehicle (15/556) or BPA (25/556) samples, suggesting that differential enhancer identity was not the primary mechanism driving changes in chromatin accessibility (Figure 3D).

**Figure 3.**
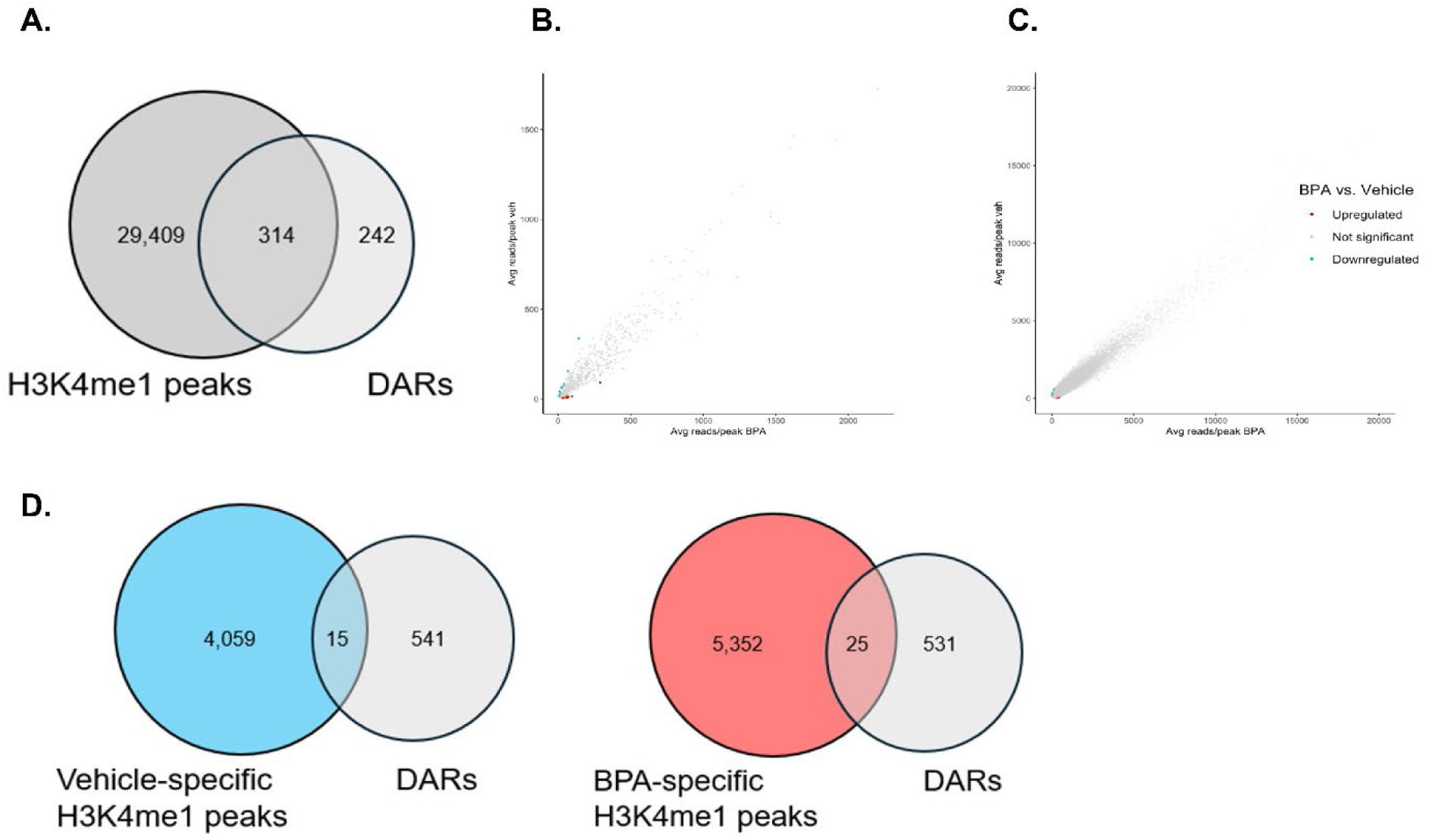
BPA drives chromatin changes in enhancer elements. CUT&Tag was performed for H3K4me1 in mammary fibroblasts. Peaks were called in each treatment group against an IgG control using MACS3 and merged using mergePeaks to obtain a union set of H3K4me1 peaks. H3K4me1 peaks (dark grey) were overlapped with the ATAC-seq DARs (light grey) using mergePeaks **(A)**. Reads were counted in DARs and normalized reads were averaged within treatment groups (n = 3) and plotted as counts in vehicle vs. BPA treated samples. Differentially marked regions for BPA (red) and vehicle (blue) were defined using │log2FC│ ≥ 1.5 and p ≤ 0.05 **(B)**. Reads were counted in H3K4me1 peaks and normalized reads were averaged within treatment groups (n = 3) and plotted as counts in vehicle vs. BPA treated samples. Differentially marked regions for BPA (red) and vehicle (blue) were defined using │log2FC│ ≥ 1.5 and FDR ≤ 0.05 **(C)**. H3K4me1 peaks called uniquely in vehicle (blue) or BPA (red) treatment groups were overlapped with DARs using mergePeaks **(D)**.

Because chromatin profiles were not shifted consistently in one direction, which could indicate a mechanism driven by distinct chromatin remodeling events, we explored the possibility that changes in chromatin accessibility at enhancers within our bulk populations could be due to changes in fibroblast heterogeneity. Interestingly, 121 out of 567 DARs (21%) were annotated to genes shown to be differentially expressed in specific mammary fibroblast subtypes, in comparison to an average of 30 out of 567 random genes (5%) from 3 independent randomly generated gene sets (Figure 4A)^41^. Enhancers that were differentially accessible in vehicle treated fibroblasts were nearby to genes associated with a progenitor-like fibroblast state, such as Sema3c, Creb5, Psd3, and Klf4 (Figure 4B). In contrast, ATAC signal in BPA treated fibroblasts was enriched in regions nearby to genes marking more differentiated fibroblasts, such as Dapk1, Cxcl12, Loxl1, and Sh3pxd2b (Figure 4C). These results suggested that in utero BPA exposure may cause an enrichment of differentiated fibroblasts in the mammary gland.

**Figure 4.**
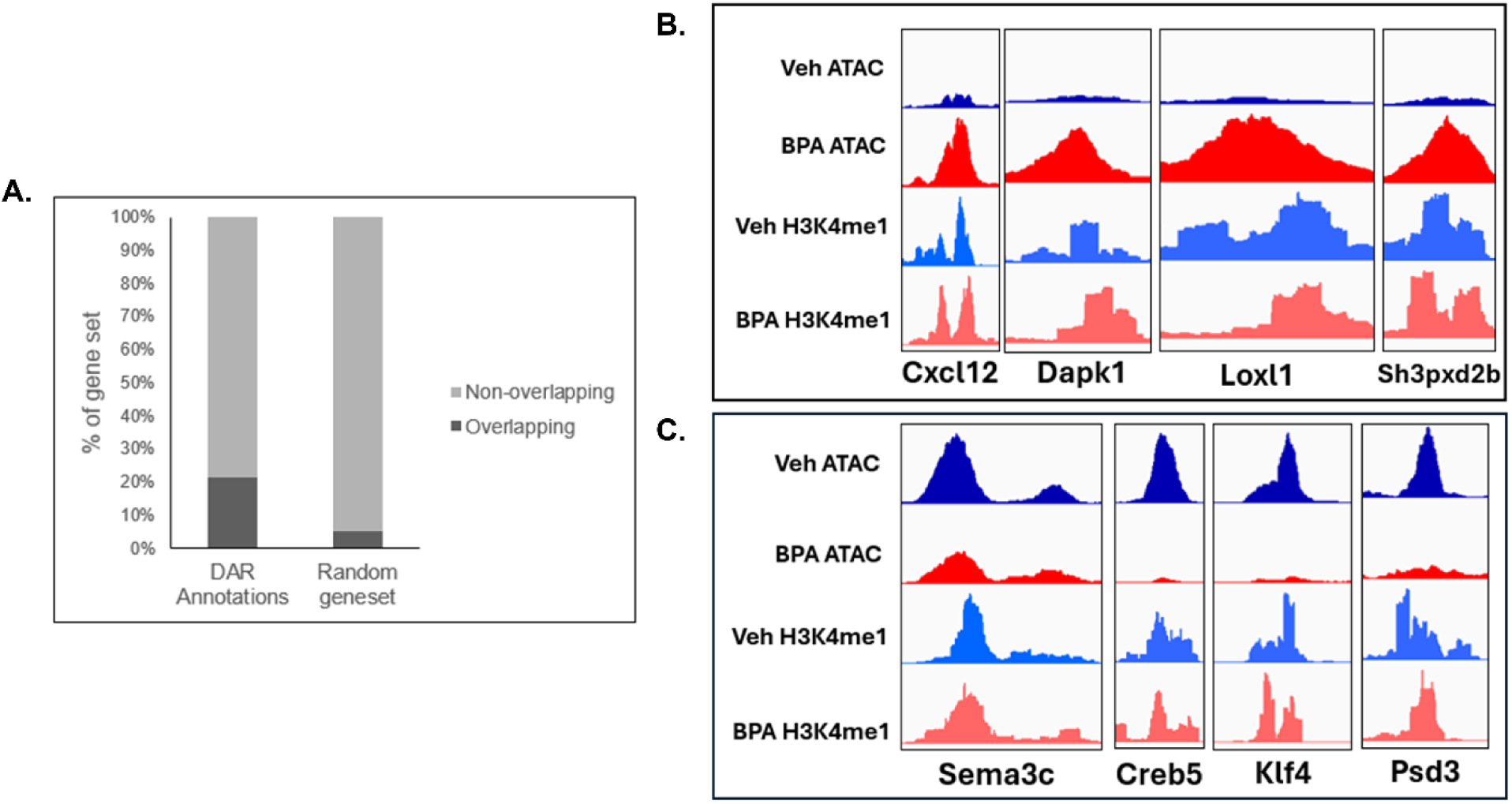
Chromatin alterations are enriched near genes associated with fibroblast differentiation. DARs were annotated to the closest gene using HOMER annotatePeaks and nearby genes were overlapped with markers of fibroblast subtypes from Yoshitake et al. The percentage of gene annotations overlapping cell type identity genes (dark grey) were compared for DARs and the average of 3 random gene sets **(A)**. IGV screenshots show chromatin accessibility (darker colors) and H3K4me1 (lighter colors) in vehicle (blue) and BPA (red) samples for genes marking a progenitor state **(B)** or a differentiated state **(C)**.

### BPA exposure affects cellular heterogeneity in the mammary gland

The proximity of differentially accessible and H3K4me1-marked regions to genes involved in cell type identity suggested that BPA exposure could reprogram fibroblast heterogeneity. To investigate this hypothesis and further explore the gene regulatory networks governing fibroblast heterogeneity, we performed single nuclei RNA/ATAC-seq at 12 weeks post-exposure. Cells were clustered using a weighted nearest neighbor (WNN) algorithm, which revealed 17 clusters that expressed common cell type markers (Supplemental Figure 1A,B). Clusters were annotated based on their expression of known cell type markers and their similarity to published datasets (Figure 5A)^46,47^. We next sought to identify changes in the cellular composition of the mammary gland following in utero BPA exposure. One of the biggest changes in percentage of cell types was a significant increase in vascular endothelial cells (Supplemental Figure 1C). This increase was not validated by flow cytometry of the mouse mammary gland (Supplemental Figure 1D), and thus vascular endothelial cells were excluded from cellular composition analyses.

**Figure 5.**
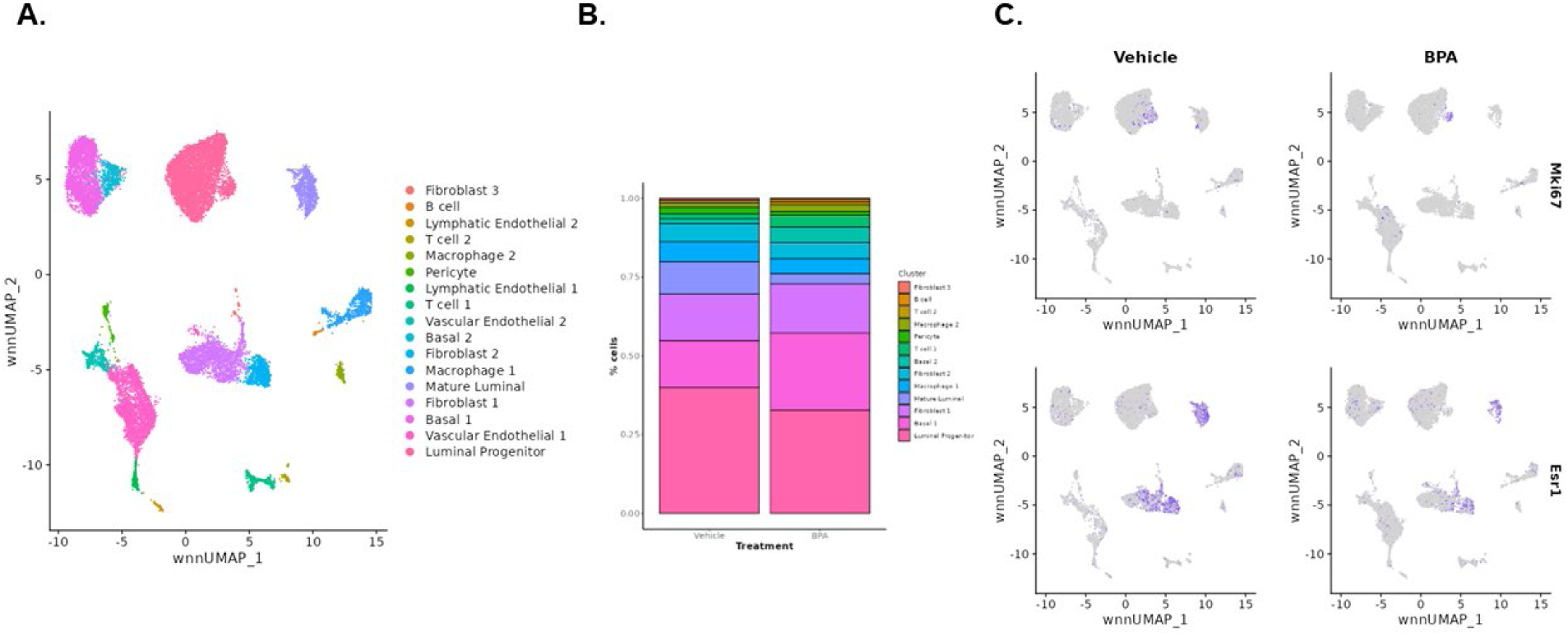
Single cell analysis recapitulates known phenotypes of BPA exposure. Single nuclei multiomics was performed using the 10X RNA/ATAC chemistry on cell pellets from adult mammary glands. Data was normalized, samples were integrated, and cells were clustered using a weighted nearest neighbor (WNN) algorithm considering both the RNA and ATAC modalities. Cell clusters were annotated using common RNA markers (A). Cell type counts were determined as a percentage of the sample for each treatment group (B). (C.) Shows dimensional reduction plots with gene expression for Mki67 or Esr1 overlaid. Cells with purple color express higher levels of the labeled marker.

Plotting the changes in cell type percentages between each treatment group revealed that BPA increased the percentage of basal cells and decreased the percentage of both HR- luminal progenitor (LP) and HR+ mature luminal cells (ML) (Figure 5B). This is in line with previous reports that BPA decreases ER positivity in mammary luminal cells^12^. This confirmed to us that our single cell sequencing accurately recapitulated the profile of mammary glands exposed in utero to BPA. There was additionally a small increase seen in the percentage of fibroblast cluster 1, while the percentage of fibroblast cluster 2 decreased with BPA exposure. We additionally showed that BPA treatment reduces the number of Mki67 positive LP cells and Esr1 positive ML cells and fibroblasts, further confirming that our multiome dataset accurately profiled BPA-exposed mammary glands (Figure 5C). While we performed differential gene expression analysis (Supplemental Files), our results were in line with documented limitations in single cell methodologies to identify differentially expressed genes (DEGs) from single samples, with significant changes in highly variable genes and more changes in clusters with dramatic differences in cell number^49^. Grouped luminal, basal, and fibroblast populations additionally shared very few DEGs with those identified in published bulk RNA-seq experiments, further confirming that single-sample gene expression analysis for scRNA-seq is unreliable (Supplemental figure 1E). Thus, we focused our analysis on differentiation and its regulators due to bulk ATAC data suggesting that BPA may reprogram fibroblast heterogeneity.

Because bulk data suggested that BPA may impact genes relating to fibroblast identity and cell type analyses showed shifts in fibroblast proportion, we subset and reclustered fibroblasts to further examine fibroblast differentiation following BPA exposure, which led to the identification of 5 subclusters of transcriptionally distinct fibroblasts (Figure 6A,B). C0 was defined by traditional fibroblast markers such as Vim and Egr1 (Figure 6B). C1 strongly expressed markers of the Pi16 signature, which has previously been shown to mark a progenitor fibroblast population in the mammary gland that is reduced with estrogen (Figure 6B, C)^41,55^. C2 expressed markers of further differentiation and ECM production, while C3 was defined by the enrichment of epithelial markers. The genes marking C4 aligned with a previously published signature of adipo-regulatory fibroblasts in the mammary gland^41,55^.

**Figure 6.**
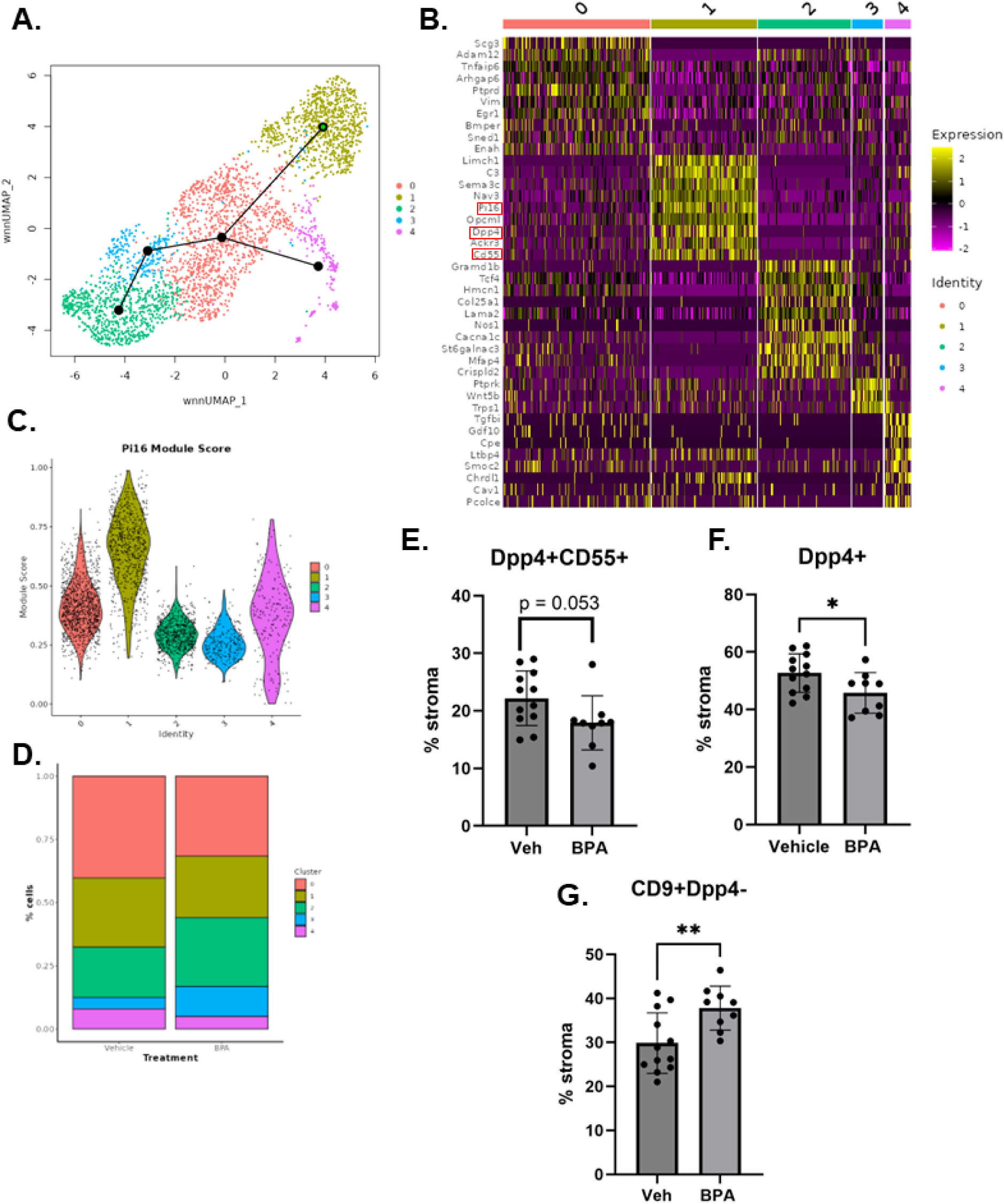
In utero BPA exposure drives fibroblast differentiation. Fibroblasts were subset and reclustered using the WNN algorithm. Five subclusters were identified and their lineage trajectory was determined using Slingshot with C1 designated as the starting point **(A)**. Transcriptional markers of each cluster were determined using FindAllMarkers on the RNA modality and visualized with a heatmap **(B)**. A module score was added to each cluster for the Pi16+ progenitor fibroblast RNA signature and visualized using a violin plot **(C)**. Cell type counts of each subcluster were visualized as a percentage of total fibroblasts for each treatment group **(D)**. Flow cytometry was used to validate fibroblast identity using CD55 and DPP4 as markers of the progenitor state and CD9 as a marker of the differentiated state. Percentage of lineage negative cells (stroma) double positive for DPP4 and CD55 **(E)**, positive for DPP4 **(F)**, and positive for CD9 **(G)** are shown.

Guided by the identification of C1 as Pi16+, trajectory analysis was performed with Slingshot, identifying a differentiation trajectory that progresses from C1 to C0, with C3 and C2 being more differentiated and C4 existing in a separate lineage derived from C0 (Figure 6A, line overlay). Cell type proportion analysis identified that the BPA-exposed sample was enriched for fibroblast subpopulations that are further along the differentiation trajectory (Figure 6D). Using scRNA-seq data from an initial single cell multiome experiment which had low quality ATAC-seq data also showed similar trends with BPA reducing the percentage of Pi16+ fibroblasts and increasing the percentage of more differentiated fibroblasts (Supplemental Figure 2A-C).

To further validate this finding, we designed a flow cytometry panel guided by our multiomics data, which identified DPP4 and CD55 as markers of Pi16+ progenitor fibroblasts (Figure 6B, boxes). Previously published scRNA-seq data showed that Cd9 is more highly expressed in Pi16 negative fibroblasts and thus we used CD9 as a marker of differentiated fibroblasts^41^. Fibroblasts were identified in this panel as the population negative for lineage (CD45-CD31-) and epithelial (CD49f-CD24-) markers. In 12-week old mice, BPA exposure decreased the percentage of DPP4+CD55+ fibroblasts (p = 0.053) and significantly decreased the percentage of total DPP4+ fibroblasts (CD55+ and CD55-). Additionally, BPA increased the percentage of CD9+DPP4-fibroblasts, further confirming that BPA reduces the percentage of progenitor fibroblasts while enriching for more differentiated fibroblasts (Figure 4E). Interestingly, the percentage of total DPP4+ fibroblasts did not change at 4 weeks of age (Supplemental Figure 2D), indicating that BPA-induced changes in fibroblast differentiation may be another phenotype that arises during puberty.

In order to explore the chromatin landscape of fibroblast subtypes and relate changes in differentiation to the results obtained from bulk ATAC-seq, we utilized the ATAC modality of the single-nuclei multiomics dataset. Qualitatively, we observed that DARs from bulk fibroblasts were differentially accessible in fibroblast subtypes and linked to gene expression from their nearest promoters (Figure 7A). Quantitatively, we identified ATAC peaks marking each fibroblast subtype and found that 37% of bulk DARs (206/556) overlapped with these regions (Figure 7B). DARs that overlapped subtype-specific peaks were enriched for AP-1 motifs, suggesting that these transcription factors are important for regulation of cell type specific enhancers that change with BPA exposure (Figure 7C). Very few DARs were identified in the analysis of fibroblast subclusters (Figure 7D), suggesting that differences in cell state were a large contributor to differential accessibility identified in our bulk ATAC-seq experiment. However, a single multiome sample has limited statistical power to identify differentially accessible peaks. Similarly to the results of our RNA analysis, very few DARs overlapped between those obtained from bulk ATAC-seq data and those obtained from grouped fibroblast clusters (Supplemental Figure 3A), confirming the limitations of differential accessibility analysis using single nuclei data.

**Figure 7.**
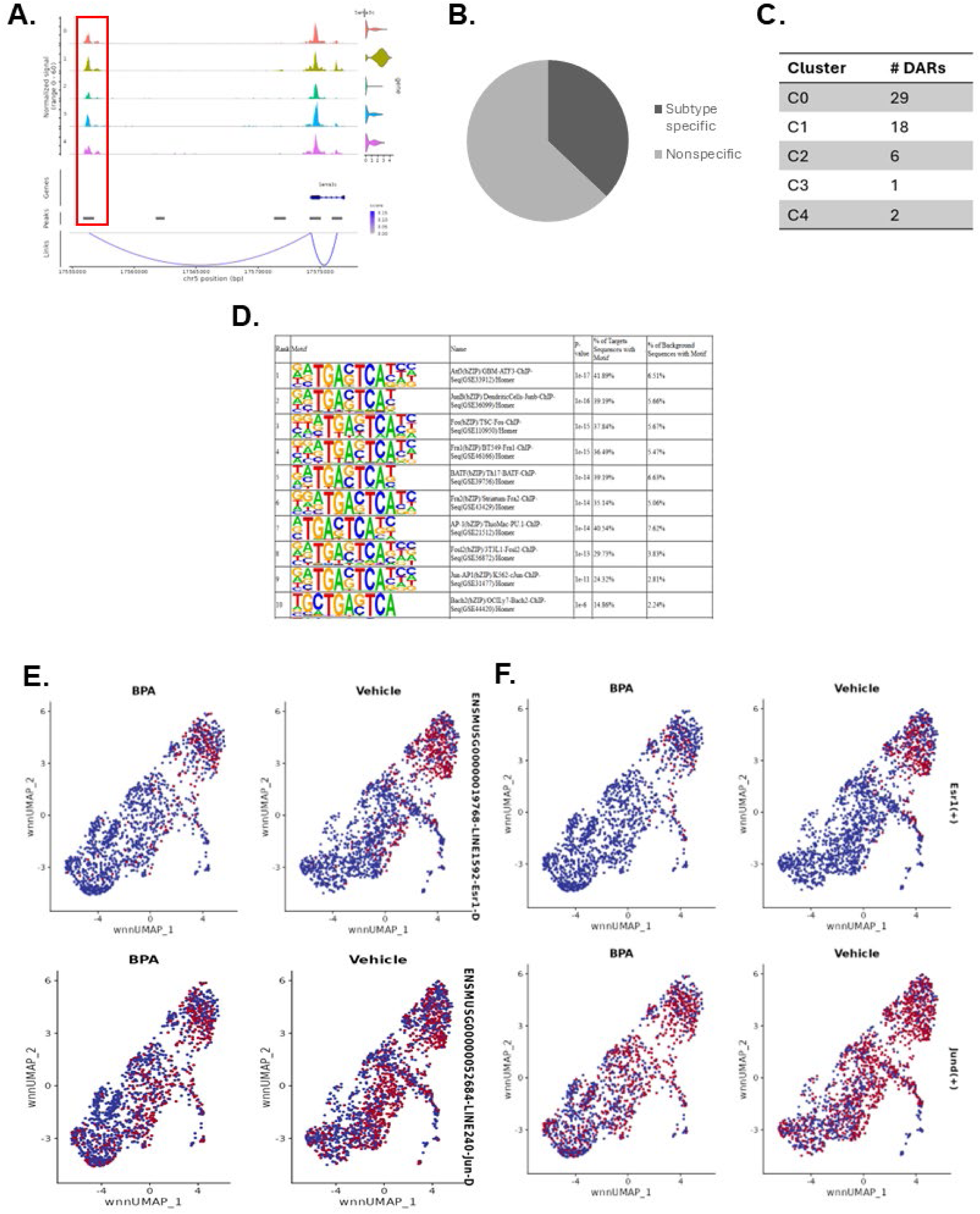
In utero BPA exposure reprograms differentiation genes associated with the AP-1 and estrogen signaling pathways. The ATAC modality of fibroblast subclusters was analyzed for differential accessibility, overlap with bulk data, motif enrichment, and gene regulatory network construction. (A.) Coverage plot showing the Sema3c locus in each fibroblast subcluster, with ATAC accessibility along the horizontal axis and corresponding RNA expression along the vertical axis. The red box highlights the Sema3c enhancer displayed in Figure 3-4. Peaks called by CellRanger are represented by gray boxes below the accessibility track and purple lines show the predicted linkages between peaks and gene promoters. (B.) Pie chart showing the proportion of differentially accessible regions from bulk ATAC-seq data in fibroblasts that overlap with subtype-specific regions identified by single cell sequencing. (C.) Table showing the number of differentially accessible regions (logFC threshold 0.5, padj < 0.05) identified for each fibroblast subcluster. (D.) HOMER results showing enriched motifs for bulk DARs that are fibroblast subtype-specific. (E.) Dimensional reduction WNN UMAP split by treatment showing the accessibility of the Esr1 (top) and Jun (bottom) motifs. Color on a blue to red scale represents the enrichment of the given motif, with cells in red having higher motif accessibility. (F.) Dimensional reduction WNN UMAP split by treatment showing the expression of the Esr1 (top) and Jund (bottom) regulatory network genes. Color on a blue to red scale represents the expression of the regulome, with cells in red having higher expression.

Rather than using differential accessibility of individual peaks, we sought to identify potential regulators of the differentiated fibroblast state altered by BPA using ChromVAR analysis to examine motif accessibility in fibroblast subtypes and how it changes with each treatment. Examining motifs that mark each subtype revealed enrichment of several transcription factors known to be involved in fibroblast differentiation in other organs, such as Klf1 and Runx1 (Supplemental Figure 3B)^56,57^. We then sought to identify motifs that may be affected by BPA exposure, which revealed several cluster-specific motifs whose accessibility is altered between BPA and vehicle treatment (Supplemental File). Interestingly, both the Esr1 and Jun motifs were significantly less accessible in BPA treatment (Figure 7E), further supporting bulk ATAC-seq data that identified AP-1 motifs as significantly enriched in DARs.

The ATAC and gene expression modalities were integrated for gene regulatory network (GRN) analysis using Pando to identify regulators of the differentiated fibroblast state. Pando identifies regulatory modules based on expression of genes and motif enrichment in nearby accessible peaks^52^. GRN analysis was limited to gene markers of fibroblast subclusters in order to identify regulatory modules that could be driving fibroblast differentiation. Module scores for the resulting GRNs were calculated and compared between vehicle and BPA treatment for each cluster to determine how BPA may regulate fibroblast differentiation (Supplemental File). In the Pi16+ C1, BPA treatment significantly reduced the expression of the genes positively regulated by Esr1 (Figure 7F). Across all clusters, BPA significantly reduced the expression of genes in the Jund positive regulon (Figure 7F). This analysis was concordant with our ChromVAR analysis, showing both accessible motifs for and their linked genes are differentially regulated in BPA treatment for both Jund and Esr1. ERα is known to be tethered by transcription factors from the AP-1 family, and thus this data suggests that the two transcription factors may synergize to govern fibroblast differentiation after BPA exposure. Overall, this supports a model whereby BPA reprograms the estrogen responsiveness of mammary fibroblasts to affect their differentiation, ultimately leading fibroblasts to adopt a more differentiated, pro-secretory state that then creates a microenvironment supporting mammary tumorigenesis.

## Discussion

Decades of work have established that in utero exposure to BPA alters mammary gland development and increases tumor susceptibility. However, the molecular mechanisms underlying these alterations and their propagation from womb to adulthood are still unclear, making it difficult to fully assess the risk of exposure and design safe alternatives. We thus aimed to determine both the timing of BPA-induced changes in transcription and identify their potential regulators. Our findings support a model where BPA acts to reprogram fibroblast differentiation in response to ovarian hormones through differential usage of cell-type specific enhancers.

Because we previously showed that pre-pubertal mice exposed in utero to BPA had no changes in collagen deposition or tissue stiffness, we first sought to investigate if this phenotype is regulated at the transcriptional level^17^. We confirmed that prior to puberty, there are no differences in collagen mRNA expression, and that collagen begins to be upregulated at 7 weeks of age. This suggests that the reprogramming by BPA is initiated in response to the mammary gland maturation signals during puberty, and even possibly to the increase in circulating ovarian hormones. This conclusion is supported by prior work in the mouse uterus showing that the effects of DES exposure manifest after a secondary exposure to endogenous estrogen^58^. Prior studies in the mammary gland also concluded that estrogen response is altered after in utero exposure to BPA^59,60^. Studies of the hepatic response to estrogen show that estrogen exposure primes genes to transcriptionally respond to estrogen later through reconfiguration of chromatin structure, providing a potential explanation for how BPA may induce these changes in the mammary gland^61^. Due to the manifestation of transcriptional changes much later than the initial exposure, we sought to identify changes in epigenetic regulation that could be driving this change.

While others have profiled the epigenome of BPA-exposed mammary glands, we separated the mammary gland into individual cell types, showing that fibroblasts are the primary target of epigenetic dysregulation in the mammary gland. Differentially accessible regions in mouse fibroblasts were found to be distal to transcription start sites and overlapped with enhancers marked by H3K4me1. However, there were very few changes in levels of H3K4me1, suggesting that enhancer identity itself is not the major driver of changes in chromatin accessibility in adulthood. One potential explanation for this observation is that BPA exposure changes how enhancers are selected in fibroblasts. Enhancer selection can be governed by binding of lineage determining transcription factors such as AP-1, and other chromatin marks such as DNA methylation or H3K27me3, that prime enhancers for binding by signal dependent transcription factors such as ERα; these marks present avenues for future investigation of the mechanism of reprogramming by BPA^62^. Prior study of epigenomic reprogramming by BPA in the liver also found very few changes in H3K4me1 in adulthood, instead showing that BPA accelerates the accumulation of H3K4me1 at sites that normally gain this mark with age^63^. However, some sites in this study did show persistent differences in H3K4me1 across different timepoints, and thus it will be critical in the future to profile the epigenome at the time of exposure to determine persistent changes in the mammary gland epigenome.

Interestingly, genes nearby to DARs were not enriched in terms relating to extracellular matrix production or remodeling, suggesting that BPA does not directly regulate collagen production. However, many DARs were annotated to genes specific to fibroblast subtypes, suggesting that these changes could be related to fibroblast heterogeneity. Single cell and flow cytometry data confirmed that BPA-exposed fibroblasts were enriched for a more differentiated state marked by the expression of ECM proteins. This is in line with prior data showing that estrogen decreases the percentage of Pi16+ progenitor fibroblasts in the mammary gland^55,64^. These changes in fibroblast heterogeneity coupled with the lack of enrichment of ECM related terms suggests that changes in collagen deposition in the mammary gland after BPA exposure may be driven by fibroblasts existing in a more differentiated, pro-secretory state.

The trajectory of fibroblast differentiation in the mammary gland is presently unknown. Interestingly, picrosirius red staining at 8 and 9 weeks of age suggests that collagen deposition decreases over time, but the magnitude of this decrease is smaller for BPA-treated mice. Similarly, our flow cytometry data shows that the percentage of DPP4+ progenitor fibroblasts increases between 4 and 12 weeks of age, which is further confirmed by a published scRNA-seq study showing that progenitor-like fibroblasts increase with age in the mammary gland^65^. These observations suggest that BPA could be affecting the transition of mammary fibroblasts between progenitor and differentiated states. Supporting this hypothesis, 37% of DARs from bulk ATAC-seq overlap with subtype-specific peaks identified in single nuclei ATAC data. In the future, it will be critical to understand how these regions change during differentiation in order to better understand how BPA reprograms the epigenome to alter fibroblast differentiation.

To begin understanding the potential regulators of the fibroblast state that may be altered by BPA, we jointly analyzed the RNA and ATAC modalities, which revealed that Esr1 and Jun regulomes are significantly altered by BPA. This analysis further supported the conclusions of our prior work, which showed that engagement of ERα is important for EDC-driven stiffness and tumor susceptibility^18,21^. AP-1 motifs were also significantly enriched in both DARs overall and in DARs that overlapped with fibroblast subtype-specific peaks. AP-1 transcription factors are known to tether nuclear hormone receptors, including ERα, and this was previously shown to be an important mechanism regulating androgen receptor activity in prostate fibroblasts^66^. The mechanism of genomic ERα signaling in mammary fibroblasts is presently unknown, but our data suggests that similarly to the prostate, it may involve tethering with AP-1 to influence gene regulation. In the future, it will be critical to understand how ERα signals in mammary fibroblasts in order to accurately determine the mechanism of transcriptional reprogramming by endocrine disruptors. Altogether, integrating this work in the context of the field of endocrine disruption supports a model where EDCs act on mesenchymal ERα to reprogram the estrogen response of mammary fibroblasts, leading to changes in fibroblast differentiation that prime the mammary microenvironment for tumorigenesis to occur.

## Supporting information

Folder of DEG lists in each cell type

Differential scATAC motif accessibility

Differential GRN enrichment

## Supporting Information

The following files are available free of charge.

Supplemental figures 1-3 (PDF)

Differential gene expression results (.zip folder of .csv)

Differential motif accessibility results (.csv)

Differential gene regulatory network results (.csv)

## Data Availability

All sequencing data generated in this study are available in the NCBI Sequencing Read Archive (SRA) under BioProject PRJNA1492094.

## AUTHOR INFORMATION

### Author Contributions

Jillian Poska: conceptualization, data curation, formal analysis, writing. Bader Albalawi: data collection and analysis – flow cytometry. Madeline Price: data collection and analysis – timecourse tissue staining. Craig J. Burd: conceptualization, funding acquisition, resources, supervision, review and editing of manuscript.

## ACKNOWLEDGMENT

This study was supported by the National Institute of Environmental Health Sciences (1R01ES032026 to Craig Burd) and Pelotonia (graduate fellowship to Jillian Poska). Single nuclei multiomic sequencing was performed in collaboration with the OSUCCC Genomics Shared Resource (subsidized by CCSG P30CA016058). The authors would like to acknowledge the members of the 10X Genomics support team for their assistance in optimizing library preparation and Adam Helton for his assistance in developing the flow cytometry antibody panel.

