## Supplemental Figures 1-3 for "In utero Bisphenol A exposure developmentally reprograms mammary gland fibroblasts"

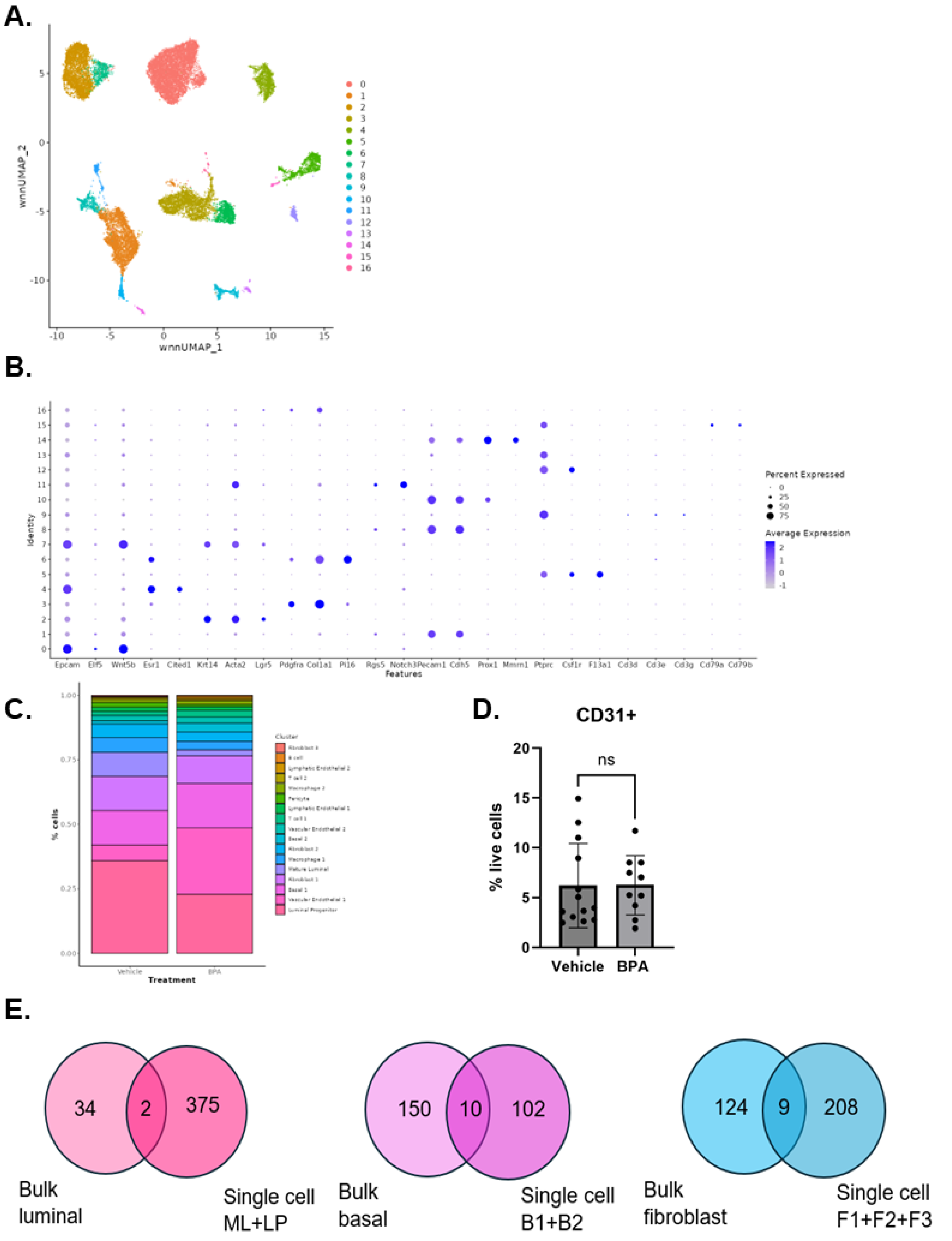


*Supplemental Figure 1. Cells were clustered using the weighted nearest neighbor algorithm with a resolution of 0.2. (A). Original WNN UMAP dimensional reduction plot showing clusters*

*labeled 0-16. (B). Dot plot demonstrating expression of luminal AV (Elf5, Wnt5b), mature luminal (Esr1, Cited1), basal (Krt14, Acta2, Lgr5), fibroblast (Pdgfra, Col1a1, Pi16), pericyte (Rgs5, Notch1), endothelial (Pecam1, Cdh5, Prox1, Mmrn1), and immune (Ptprc, Csf1r, F13a1, Cd3, and Cd79) markers. Size of the dot corresponds the percentage of cells in the cluster expressing that marker, while color on a gray to purple scale corresponds with the expression level. (C). Bar plot quantifying the percentage of each cell type per sample for each treatment. Vascular endothelial cells are included. (D.) Bar plots showing the percentage of CD31+ vascular endothelial cells by flow cytometry. Each dot represents one mouse. (E.) Venn diagrams showing the overlap between differentially expressed genes determined by bulk or single-nuclei RNA-seq for luminal epithelial cells, basal epithelial cells, and fibroblasts.*

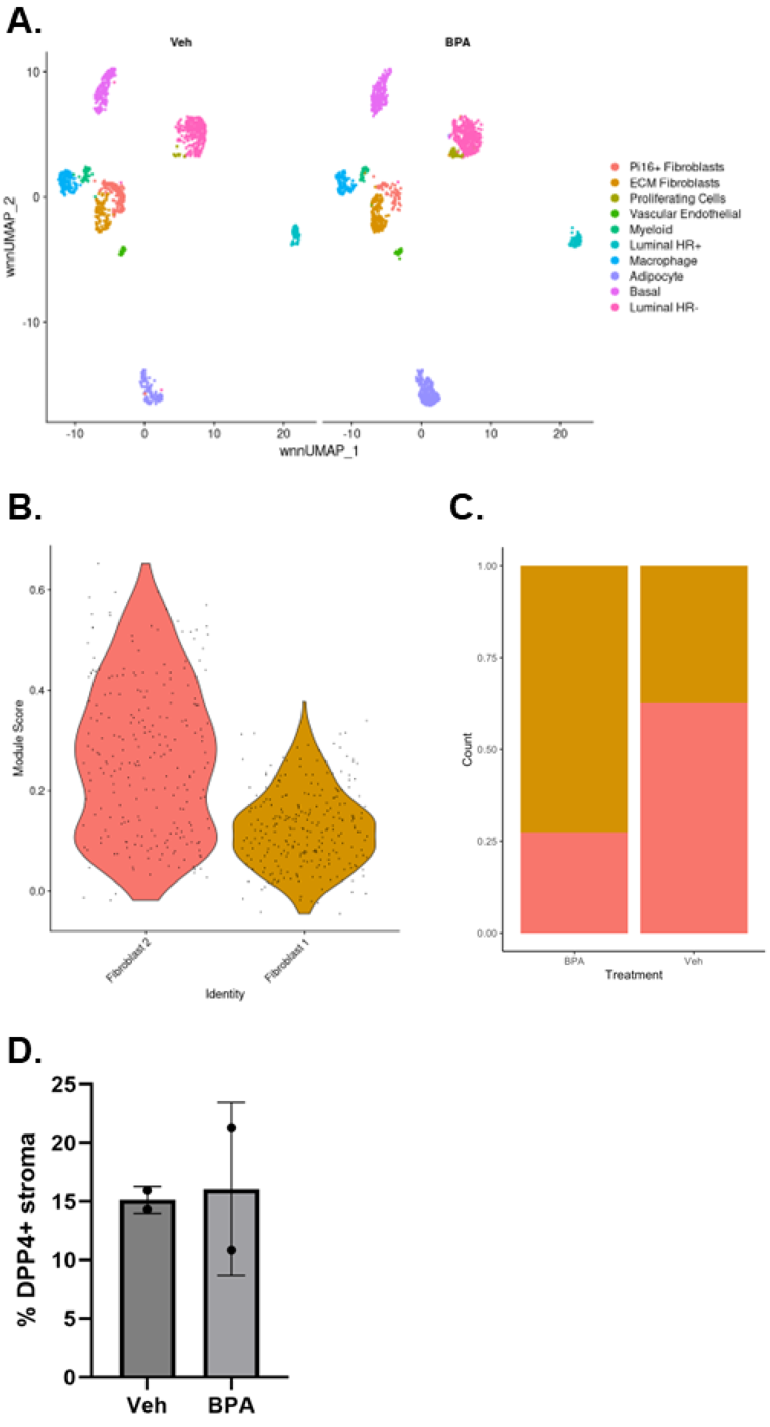


*Supplemental Figure 2. An initial single nuclei RNA/ATAC-seq experiment with nuclei isolated from frozen tissue was analyzed for cell type proportions. (A.) Dimensional reduction plot using the WNN UMAP reduction showing each identified cell type split by treatment to visualize changes in cell type proportion with each treatment. (B.) Violin plot showing the expression of the Pi16+ fibroblast gene signature in each fibroblast cluster. (C.) Bar plots showing the percentage of each fibroblast type as a proportion of total fibroblasts for each treatment. (D.)*

*Bar plots show the percentage of lineage negative cells (stroma) positive for DPP4 in 4 week old mice as determined by flow cytometry. Each dot represents one mouse.*

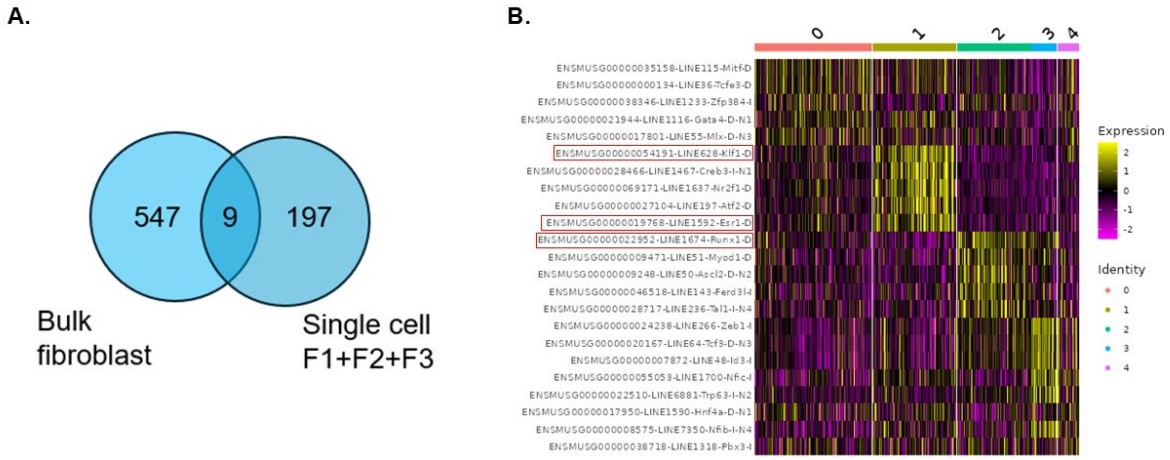


*Supplemental Figure 3. The ATAC modality of the single cell data was analyzed for differences in chromatin accessibility and motif enrichment in each fibroblast subpopulation. (A.) Venn diagrams showing the overlap between differentially accessible determined by bulk or single-nuclei ATAC-seq for fibroblasts. (B.) Heatmap showing the enrichment of motifs identified by ChromVAR in fibroblast subtypes. Yellow color corresponds to higher motif accessibility. Red boxes highlight motifs of interest (Esr1) and those previously reported to be important for fibroblast differentiation (Klf1, Runx1).*
